# Adaptation of the binding domain of *Lactobacillus acidophilus* S-layer protein as a molecular tag for affinity chromatography development

**DOI:** 10.1101/2022.12.24.521862

**Authors:** Emanuel J. Muruaga, Paula J. Uriza, Gonzalo A. K. Eckert, María V. Pepe, Cecilia M. Duarte, Mara S. Roset, Gabriel Briones

## Abstract

The SLAP_TAG_ is a novel molecular TAG derived from a protein domain present in the sequence of *Lactobacillus acidophilus* SlpA (SlpA^284–444^). Proteins from different biological sources, with different molecular weights or biochemical functions, can be fused in frame to the SLAP_TAG_ and efficiently purified by the specific binding to a bacterial-derived chromatographic matrix named here Bio-Matrix (BM). Different binding and elution conditions were evaluated to set an optimized protocol for the SLAP_TAG_-based affinity chromatography (SAC). The binding equilibrium between SLAP_TAG_ and BM was reached after a few minutes at 4°C, being the apparent dissociation constant (K_D_) of 4.3 µM, a value which is similar to different Kd determined for other S-layer proteins and their respective bacterial cell walls. A reporter protein was generated (H_6_-GFP-SLAP_TAG_) to compare the efficiency of the SAC against a commercial system based on a Ni^2+^-charged agarose matrix, observing no differences in the H_6_-GFP-SLAP_TAG_ purification performance. The stability and reusability of the BM were evaluated, and it was determined that the matrix was stable for more than a year, being possible to reuse it five times without a significant loss in the efficiency for protein purification. Alternatively, we explored the recovery of bound SLAP-tagged proteins by proteolysis using the SLAP_ASE_ (a SLAP-tagged version of the HRV-3c protease) that released a tag-less GFP (SLAP_TAG_-less). Additionally, iron nanoparticles were linked to the BM and the resulting BM_mag_ was successfully adapted for a magnetic SAC, a technique that can be potentially applied for high-throughput-out protein production and purification.

## Introduction

Proteins are biopolymers formed by a particular amino acid sequence that determines a given atomic spatial distribution, also known as protein conformation. Far from being a static structure, proteins behave as nano-machines performing precise molecular activities due to internal movements of the protein parts or protein domains ^1^. In addition, proteins can interact intramolecularly or intermolecularly with different macromolecules such as proteins, DNA, RNA, polysaccharides, or small compounds ^2^.

Proteins perform vital functions in life. For instance, some proteins degrade aliments to produce essential nutrients (like digestive enzymes), others transport critical compounds for supporting life (like hemoglobin that transports oxygen to the cells), or shape the cellular structure (like actin, tubulin, or keratin), or function as hormones (like insulin), or defense the organism against pathogens (like the antibodies) ^2^.

In addition to the multiplicity of existent polypeptides in nature, molecular biology and biotechnology have generated a myriad of chimeric proteins by combining different protein domains ^3^. These artificial proteins can be used as therapeutic tools to treat cancer, autoimmunity, or different medical conditions ^4^.

However, to gain value in therapy, in biochemical, industrial, or medical applications proteins require a highly purified, stable, and concentrated state. This state is reached by a series of steps in the process of separating the desired protein from the rest of all other proteins of a sample, free of contaminants, and preserving its biological activity. The purification process exploits differences in size, charge, hydrophobicity, ligand binding affinity, or specific sequences^5^ ^6^ ^7^. Several fractionations o chromatographic steps can be combined for an efficient enrichment or purification of a particular protein.

Affinity chromatography is a special type of liquid chromatography that exploits the existence of natural ^8^ or artificial ^9^ affinities between two moieties: the molecular target (or tag) and its ligand (the affinity ligand) which is usually immobilized onto a chromatographic stationary phase to generate a chromatography matrix. Thus, proteins of interest can be fused in-frame to different molecular tags (such as His-6X, GST, MBP, etc.) expressed and purified ideally in a single step from a complex mixture of proteins.

Since it was developed by Starkestein in 1910 ^10^, affinity chromatography was gaining popularity and centrality for many industrial processes such as the purification of proteins for diagnosis, research, and therapeutic purposes ^11^. Regulatory requirements for the purity and quality of the proteins vary greatly depending on the area of application. For instance, bio-products can be used with little purification for industrial use. Also, the recombinant protein produced for research, diagnosis, or non-clinical purposes has a less stringent regulatory approval process than proteins designed as biopharmaceuticals.

The most common strategy for affinity chromatography is the ON/OFF format. In the “ON” phase, a biological sample containing the protein of interest (that can be fused-in-frame to a molecular tag) is formulated in a specific application buffer that will favor the binding process to a ligand. Then, the sample is placed in contact with a chromatography material (or chromatography matrix) that has associated the affinity ligand that consequently will recognize and retains the tagged protein. After the washes in the “OFF” phase, an elution buffer is passed or incubated to release the tagged protein by changing the pH, the ionic strength that modifies the affinity of the interaction (non-specific elution) or the addition of the free ligand that will out-compete the retained tagged protein (bio-specific elution)^11^.

The simplicity and specificity of affinity chromatography have made this technique central for the purification of biomolecules and biopharmaceuticals.

Here, we explored the adaptation of a protein domain present in the carboxy terminus of the *Lactobacillus acidophilus* protein SlpA as a molecular tag. *In silico* analysis of this region (SlpA^284–444^) allowed us to identify a tandem of two copies of the 60-aminoacid protein domain named SLP-A (pfam03217) which is necessary for the association (Fig. Supp. 1). Here, this dual SLP-A domain was named SLAP_TAG,_ and the adaptation as a molecular tag was evaluated. Recently, we have characterized the binding properties of the SLAP_TAG_ and characterized its association with the cell wall of live *Lactobacillus* for vaccine purposes ^12^. Thus, the SLAP_TAG_ was fused in frame to a chimeric antigen derived from the Shiga toxin-producing *Escherichia coli* (STEC) formed by the peptides EspA^36–192^, Intimin^653–953^ and Tir^240–378^ (or EIT). The resulting chimeric antigen (EIT-SLAP_TAG_) recombinantly expressed and purified was able to associate with the bacterial cell wall of *L. acidophilus*, a process that we named decoration. Thus, EIT-decorated *L. acidophilus* after oral administration was able to deliver the EIT antigen to the intestinal mucosa eliciting a protective immune response that controls an experimental STEC infection in mice ^12^. Remarkably, the decoration process does not modify genetically the *Lactobacillus* genome preserving its GRAS (Generally Recognized as Safe) status, a trait that is important for vaccine purposes.

Here, we explore the adaptation of the SLAP_TAG_ for the development of novel affinity chromatography, the SLAP affinity chromatography (SAC), and a comprehensive protocol is presented.

## Results

### SLAP_TAG_ binds rapidly and efficiently to the Bio-matrix (BM)

As described in the Methods section the SLAP_TAG_ (Fig. Supp.1) was fused in-frame to the carboxy-terminus of the green fluorescent protein (GFP) to generate the chimeric protein (H_6_)-GFP-SLAP_TAG_ adapted here as a SLAP_TAG_ reporter protein. As shown in Fig. 1A the structure of the chimeric protein (H_6_)-GFP-SLAP_TAG_ was predicted based on the alphaFold-2 method ^13^. The beta-barrel corresponding to GFP and the two globular domains corresponding to SLAP_TAG_ were predicted with high performance and as expected, linkers and 6xHis-tag, which are flexible regions, showed low prediction values.

**Figure 1:**
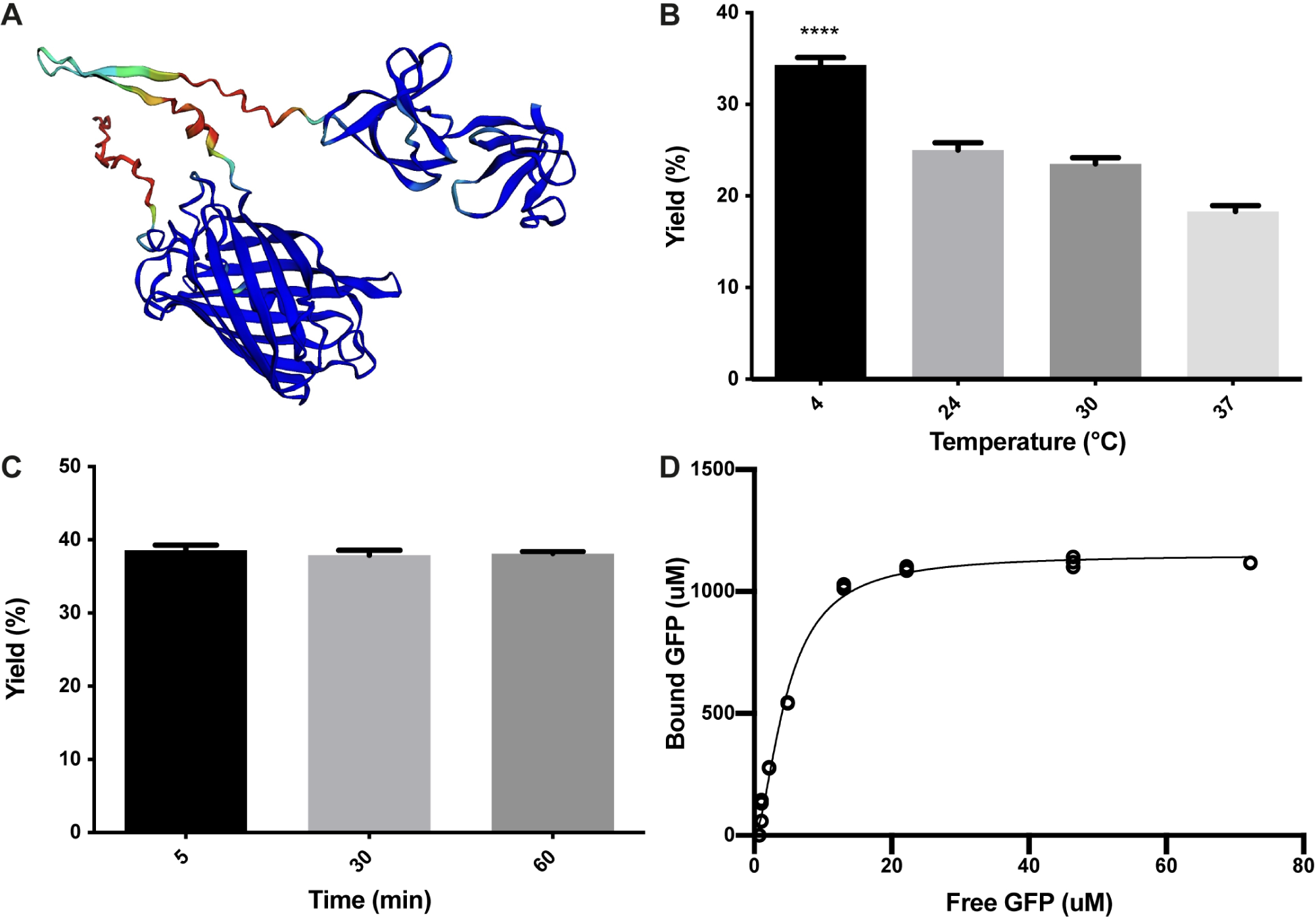
**Characterization of (H_6_)-GFP-SLAP_TAG_ binding to Bio-matrix.** (A) The AlphaFold2 model of (H_6_)-GFP-SLAP_TAG_ fusion protein. AlphaFold2 predicted structure was automatically colored by the pLDDT confidence measure. High accuracy is colored in blue while low accuracy is in red. Although SLAP_TAG_ has not been crystallized, structure prediction showed a good performance. (B) Figure compares (H_6_)-GFP-SLAP_TAG_ purification yield of SLAP_TAG_-based affinity chromatography for different temperatures of the binding process. (C) Figure compares (H_6_)-GFP-SLAP_TAG_ purification yield of SLAP_TAG_-based affinity chromatography for different times of binding incubation. (D) Adsorption isotherm of (H_6_)-GFP-SLAP_TAG_ onto Bio-Matrix. Asterisk (****) denotes significant differences using the ANOVA method, Bonferroni test (p < 0.0001).

As a working chromatographic matrix, a culture of *Bacillus subtilis* natto was processed as described in the Methods section to generate a bacterial-derived affinity chromatography matrix, named here Bio-Matrix (BM). Interestingly, although *B. subtilis* has no S-layer, we have demonstrated previously that this bacterium can be externally covered with SLAP-tagged proteins to generate a recombinant S-layer on its bacterial cell wall, a process that we called decoration ^12^. Decoration of *B. subtilis* with SLAP-tagged proteins is possible because teichoic acid and lipoteichoic acid (which are the molecules responsible for the SLAP_TAG_ association) have the same chemical composition as *Lactobacillus acidophilus*. Interestingly, in *B. subtilis* and *L. acidophilus,* teichoic acid and lipoteichoic acid are distributed homogeneously on their cell wall.

To characterize the binding properties of the SLAP_TAG_, the reporter protein (H_6_)-GFP-SLAP_TAG_, was recombinantly expressed in *Escherichia coli*, purified by affinity chromatography mediated by its Hisx6 tag (H_6_), and further incubated with BM under a variety of conditions. Initially, as shown in Fig. 1B, the optimal binding temperature of (H_6_)-GFP-SLAP_TAG_ to the BM was evaluated, finding that the best binding efficiency was reached when the incubation was performed at 4°C. To get insights into the SLAP_TAG_ association dynamics, (H_6_)-GFP-SLAP_TAG_ was incubated for 5, 30, and 60 minutes with BM, and as shown in Fig. 1C, the maximal binding of (H_6_)-GFP-SLAP_TAG_ to BM was reached very rapidly (already at 5 minutes), showing no significant increments in its association at longer time points. These results indicated that the SLAP_TAG_ binds very rapidly and with an apparent high affinity to the BM. To quantify the affinity of the interaction between the SLAP_TAG_ with the BM, an adsorption isotherm was performed to determine the apparent equilibrium dissociation constant as described in the Methods section. As shown in Fig. 1D the apparent K_D_ was estimated as 4.7 µM. Interestingly, this dissociation constant value was in the same order as other K_D_ described for different microbial S-layer proteins with their respective bacterial cell walls ^14–16^. In addition, the maximum adsorption capacity (Bmax) of BM was estimated as 1.152 mmol of (H_6_)-GFP-SLAP_TAG_ or 53.9 mg of protein per milliliter of BM or 1.03 g of SLAP_TAG_ protein/g of dry BM.

### Elution of (H_6_)-GFP-SLAP_TAG_ protein can be performed with different buffers

As it was mentioned, the SLAP_TAG_ contains the protein region responsible for the association of the protein SlpA to the *L. acidophilus* cell wall. As reported, *L. acidophilus* SlpA has the natural ability to self-assemble on the bacterial surface to generate a proteinaceous layer named S-layer ^17^, a highly ordered wall structure that functions as a protective barrier against bacteriophages, resistance to low pH and proteases, and bacterial adhesion. It is well characterized that removal of the S-layer can be performed efficiently by the addition of chaotropic agents like LiCl, a compound that disrupts hydrogen bonds leading to a partial denaturation of proteins and the consequent detachment of the S-layer ^17^. To study (H_6_)-GFP-SLAP_TAG_ elution, LiCl solution was selected as the positive control. As shown in Fig. 2A while PBS (137 mM NaCl, 2.7 mM KCl, 8 mM Na2HPO4, and 2 mM KH2PO4) does not affect elution, the detachment of BM-associated (H_6_)-GFP-SLAP_TAG_ was achieved with high efficiency with a 2M LiCl solution at room temperature, in a short lapse of five minutes (Fig. 2B). As it was mentioned above, lithium solutions have deleterious effects on proteins, which is an undesired effect for protein purification, especially for enzyme purification. Therefore, different milder alternatives of buffers were explored to elute SLAP_TAG_-tagged proteins, conditions that can preserve the protein structure and therefore their function. As shown in Fig. 2C, a series of sugars (monosaccharide and disaccharides) described previously ^18^ were tested for the elution of the reporter SLAP observing a partial elution efficiency when compared with the LiCl elution control (Fig. 2C). Next, the cationic nature of the SLAP_TAG_ at neutral pH was considered.

**Figure 2:**
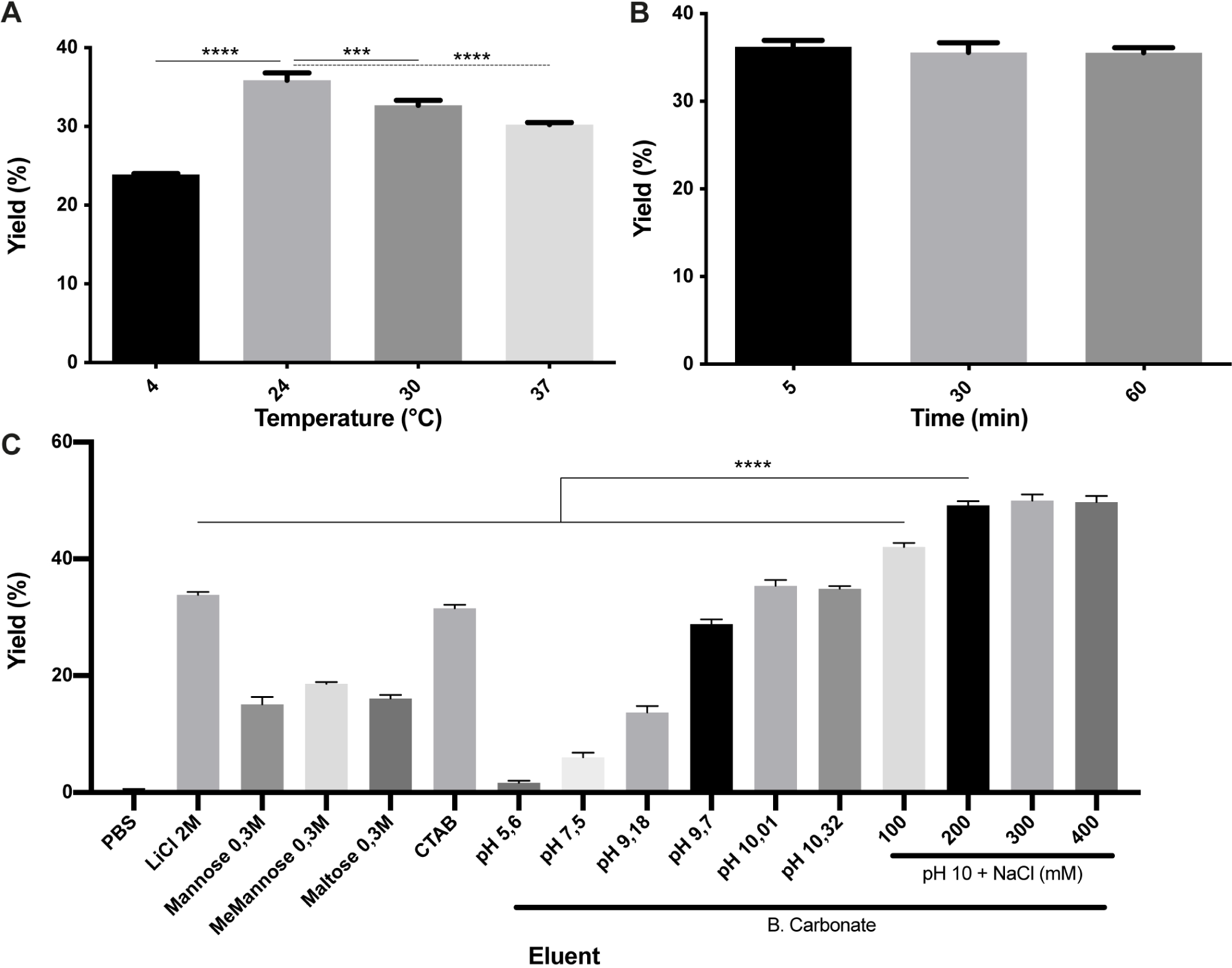
**Characterization of (H_6_)-GFP-SLAP_TAG_ elution.** (A) Figure compares (H_6_)-GFP-SLAP_TAG_ purification yield of SLAP_TAG_-based affinity chromatography for different temperatures of elution. (B) Figure compares (H_6_)-GFP-SLAP_TAG_ purification yield of SLAP_TAG_-based affinity chromatography for different times of elution incubation. (C) Figure compares (H_6_)-GFP-SLAP_TAG_ purification yield of SLAP_TAG_-based affinity chromatography system for different eluents. The asterisk denotes significant differences using the ANOVA method, the Bonferroni test (*** p = 0,0003; **** p < 0.0001).

Taking into consideration the importance of this positive charge in the interaction of SLAP_TAG_ with the membrane-bound compound teichoic acid, a set of cationic compounds was evaluated for the elution. As shown in Fig. 2C, 0.3M of CTAB was able to elute (H_6_)-GFP-SLAP_TAG_ with similar efficiency as the LiCl solutions. Although CTAB was very efficient in this step, we found that this compound was difficult to remove downstream from the elution fraction. Then, considering that SLAP_TAG_ has a theoretical isoelectric point (pI) value of 9.92, we explored if the modification of pH can be adapted as an elution method. As shown in Fig. 2C, when the pH of the bicarbonate buffer was close to the theoretical SLAP_TAG_ isoelectric point (pI), (H_6_)-GFP-SLAP_TAG_ was efficiently eluted from BM. In addition, after establishing the bicarbonate buffer at pH 10 as an elution buffer, we explore if the addition of NaCl can improve the elution process. As shown in Fig. 2C, the addition of 200 mM of NaCl was able to maximize the yield of recovery of (H_6_)-GFP-SLAP_TAG_ in the eluate. Interestingly, 200 mM of NaCl is the same concentration present in the binding/washing buffer that has no effect in detaching SLAP-tagged proteins.

### The optimized SAC protocol allowed the efficient purification of proteins with similar efficiency to the high-performance Ni^2+^-charged agarose matrix (IMAC)

With all the experimental information obtained, an optimized SAC protocol for the purification of SLAP-tagged proteins was established as shown in Fig. 3A. Remarkably, the entire process of protein purification can be performed in 15 minutes. To have a direct observation of the purification process of (H_6_)-GFP-SLAP_TAG_, the BM was fixed and immobilized on coverslips at different steps of the process of protein purification to be observed by confocal microscopy. As shown in Fig. 3B, before the incubation of BM with (H_6_)-GFP-SLAP_TAG_, only a red native autofluorescence can be observed from fixed *Bacillus subtilis* natto cells present in BM. Remarkably, after the incubation of BM with (H_6_)-GFP-SLAP_TAG,_ it was possible to detect the adhesion of the reporter protein to the BM forming a recombinant fluorescent S-layer that covers completely the bacterial surface (Fig. 3C). As expected, after the addition of the elution buffer the (H_6_)-GFP-SLAP_TAG_ was completely removed from the bacterial surface (Fig. 3D). In addition, the whole process of purification can also be monitored by direct observation under UV light (Fig. 3E)

**Figure 3:**
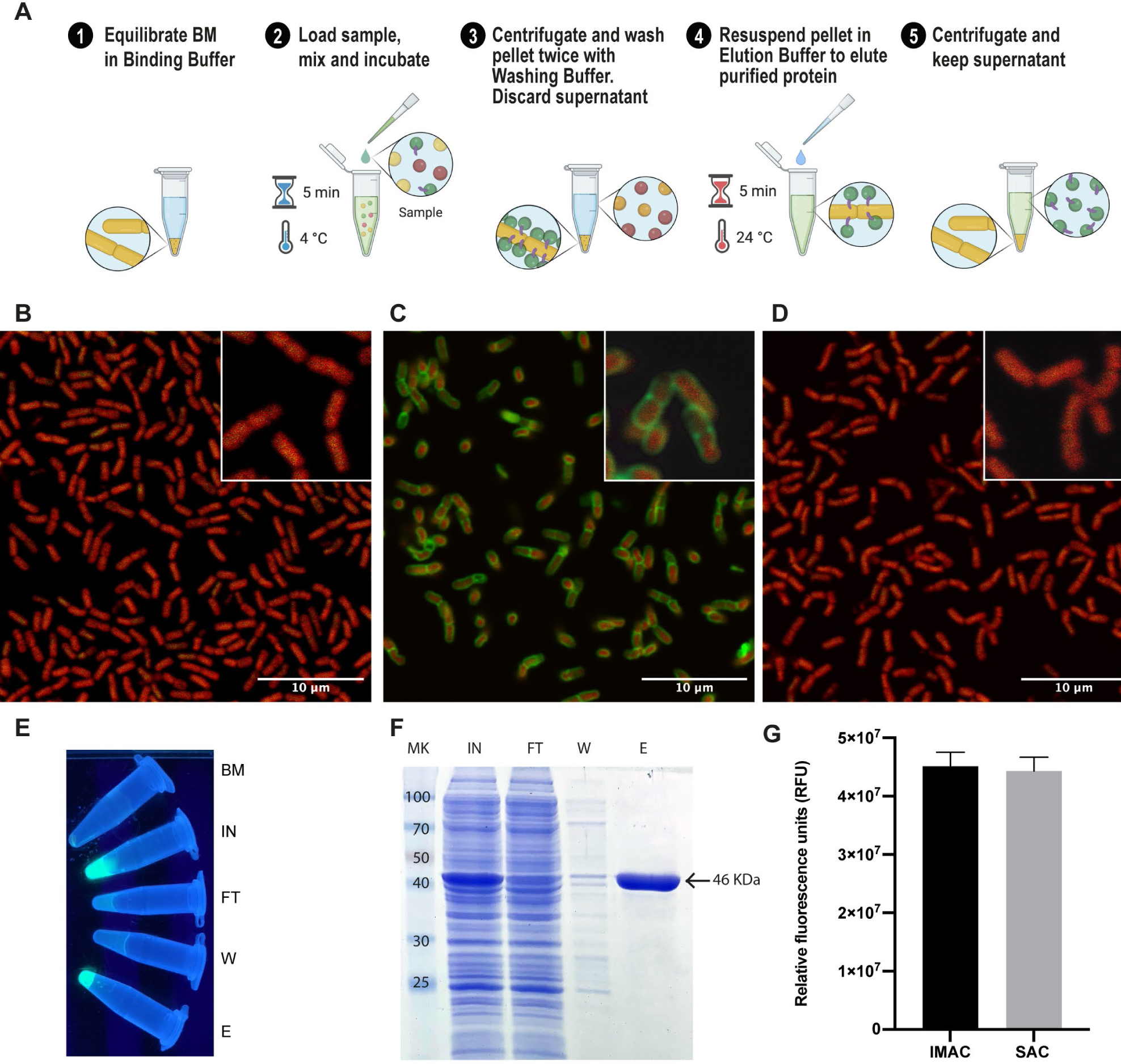
**Optimized protocol for (H_6_)-GFP-SLAP_TAG_ purification using the Bio-matrix and comparison with the high-performance Ni^2+^-charged agarose matrix.** (A) Graphical description of the SLAP_TAG_-based affinity chromatography protocol. (B) Confocal microscopy of the Bio-Matrix. *Bacillus subtilis* red native autofluorescence is observed. (C) Confocal microscopy of (H_6_)-GFP-SLAP_TAG_ bound to the Bio-Matrix. GFP is visualized on the *Bacillus* surface. (D) Confocal microscopy of the Bio-Matrix after elution of (H_6_)-GFP-SLAP_TAG_. (E) Tubes containing purification fractions seen under UV light. GFP input fluorescence is recovered in the eluate. (F) Coomassie blue staining analysis of (H_6_)-GFP-SLAP_TAG_ scaled purification. BM = Bio-Matrix; MK = protein marker (kDa); IN = input; FT = flowthrough; W = wash; E = elution. (G) Immobilized metal affinity chromatography (IMAC) and SLAP_TAG_-based affinity chromatography systems are compared in their capacity to purify (H_6_)-GFP-SLAP_TAG_. Relative fluorescence of total (H_6_)-GFP-SLAP_TAG_ recovered in elution fraction is shown.

Since SAC was efficient in protein purification, we compared this new technique against an established affinity purification protocol. For this, we took advantage of (H_6_)-GFP-SLAP_TAG_ also has a His-tag that can be purified by a Metal-Affinity-Chromatography or IMAC. As described in Materials and Methods, a bacterial lysate (H_6_)-GFP-SLAP_TAG_ was divided into two fractions and both protocols were performed accordingly in parallel, confirming that SAC optimized protocol was able to purify (H_6_)-GFP-SLAP_TAG_ with high efficiency (Table Supp. 1 and Fig. 3F) and with a similar yield to the one obtained with the commercial IMAC (Fig. 3G).

### Proteins of different biological sources, molecular weights, biochemical functions, or expressed by procaryotic or eukaryotic expression systems can be efficiently purified by the SAC

To evaluate if SLAP_TAG_ and SAC can be adapted as a universal protein purification system, a set of the selected proteins were fused to the SLAP_TAG_ (Fig. 4).

**Figure 4:**
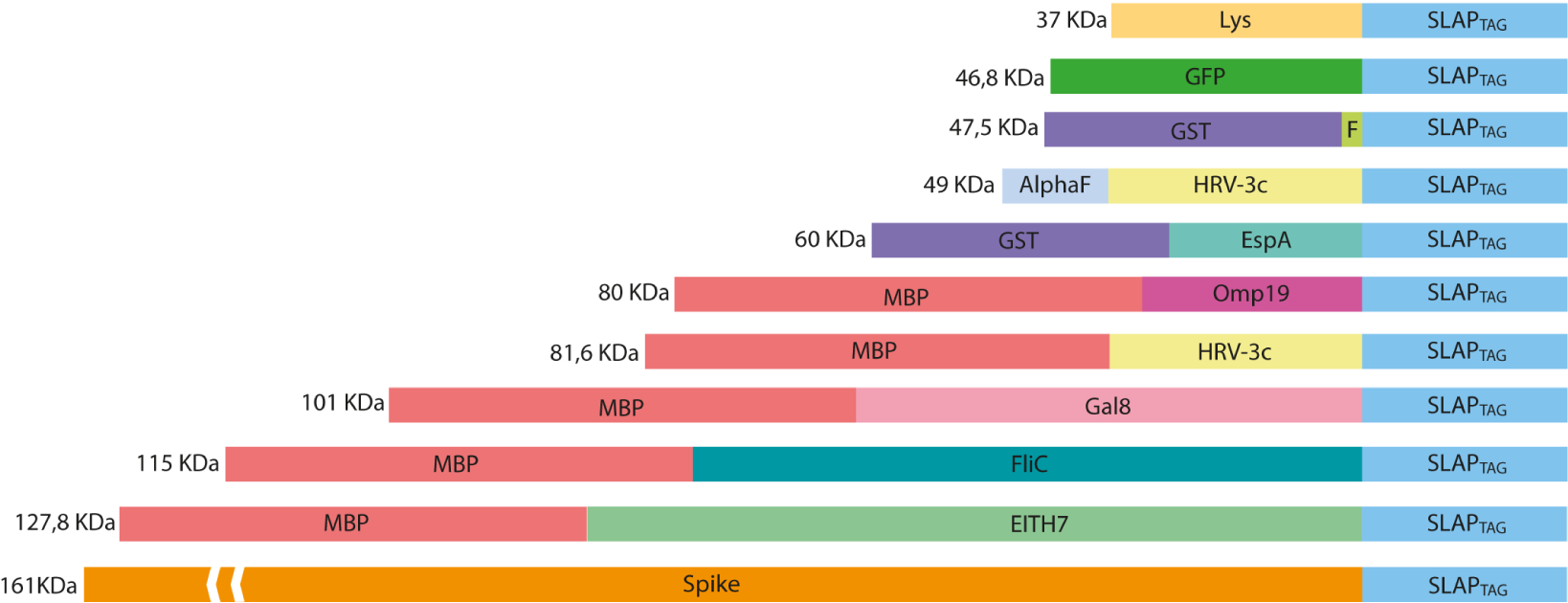
**Representation of SLAP-tagged recombinant protein structure.** Lys = bacteriophage T4 lysozyme; GFP = *Aequorea victoria* green fluorescent protein; GST = glutathione-s-transferase; HRV-3c = human Rhinovirus 3C Protease; EspA = *E. coli* EspA protein; Omp19 = *B. abortus* Omp19 protein; MBP = maltose-binding protein; Gal8 = mouse Galectin-8; EITH7 = EspA, Intimin and Tir fusion protein from the Shiga-toxin producing *E. coli*; Spike = SARS-CoV2 Spike protein.

Thus, bacterial proteins like *Salmonella* flagellin (Fig.5A), the STEC proteins, EspA (Fig. 5B), the chimeric EIT (Fig. 5C), the *B. abortus* Omp19 (Fig. 5D), the bacteriophage protein T7-lysozyme (Fig. 5E), the human viral proteins Rhinovirus 3C Protease (Fig. 5E), the mouse Galectin-8 (Fig. 5G) and the commercial molecular tag GST (Fig. 5H), were fused in frame with the SLAP_TAG_.

**Figure 5:**
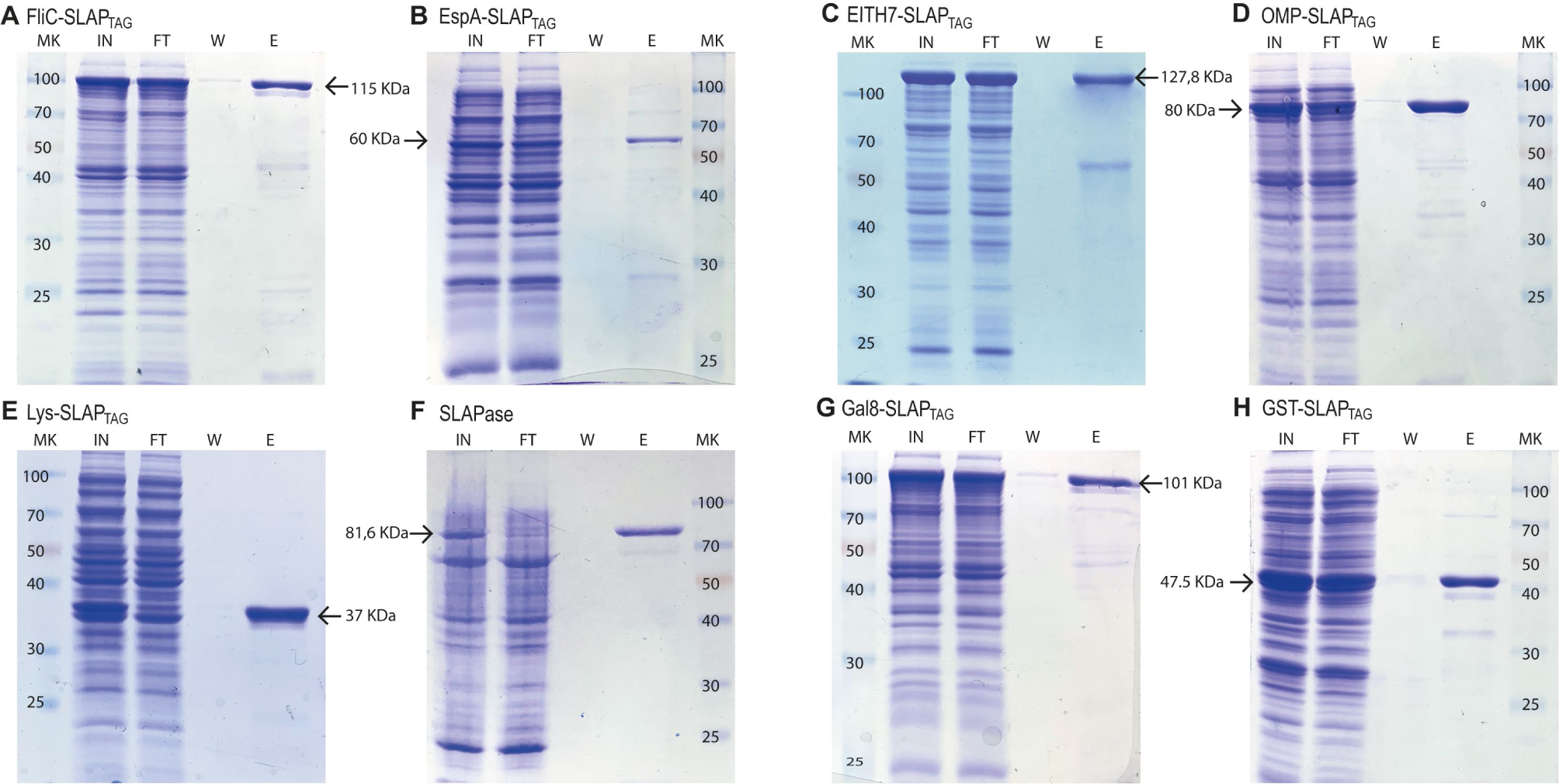
**Analysis of different SLAP-tagged protein purifications expressed in *E. coli*.** Coomassie Blue stained SDS-PAGE of purification fractions of SLAP_TAG_-based affinity chromatography for (A) FliC-SLAP_TAG_ (B) EspA-SLAP_TAG_ (C) EITH7-SLAP_TAG_ (D) Omp19-SLAP_TAG_ (E) Lys-SLAP_TAG_ (F) SLAP_ASE_ (human rhinovirus 3c fused to SLAP_TAG_) (G) Gal8-SLAP_TAG_ (H) GST-SLAP_TAG_. MK = protein marker (kDa); IN = input; FT = flowthrough; W = wash; E = elution.

As shown in Fig. 5, all the selected proteins were purified efficiently with SAC. In addition to the bacterial expression system, different protein expressions systems (yeast and mammalian cells) were also evaluated (Fig. 6). As shown in Fig. 6A the viral HRV 3C protease fused to the SLAP_TAG_ was expressed and purified from *Pichia pastoris* supernatant. Also, the SARS-CoV2 SPIKE fused to the SLAP_TAG_ was purified from the supernatant of transfected HEK293 (Fig. 6B). These results confirmed that the SLAP_TAG_ can be widely adopted for affinity chromatography purification.

**Figure 6:**
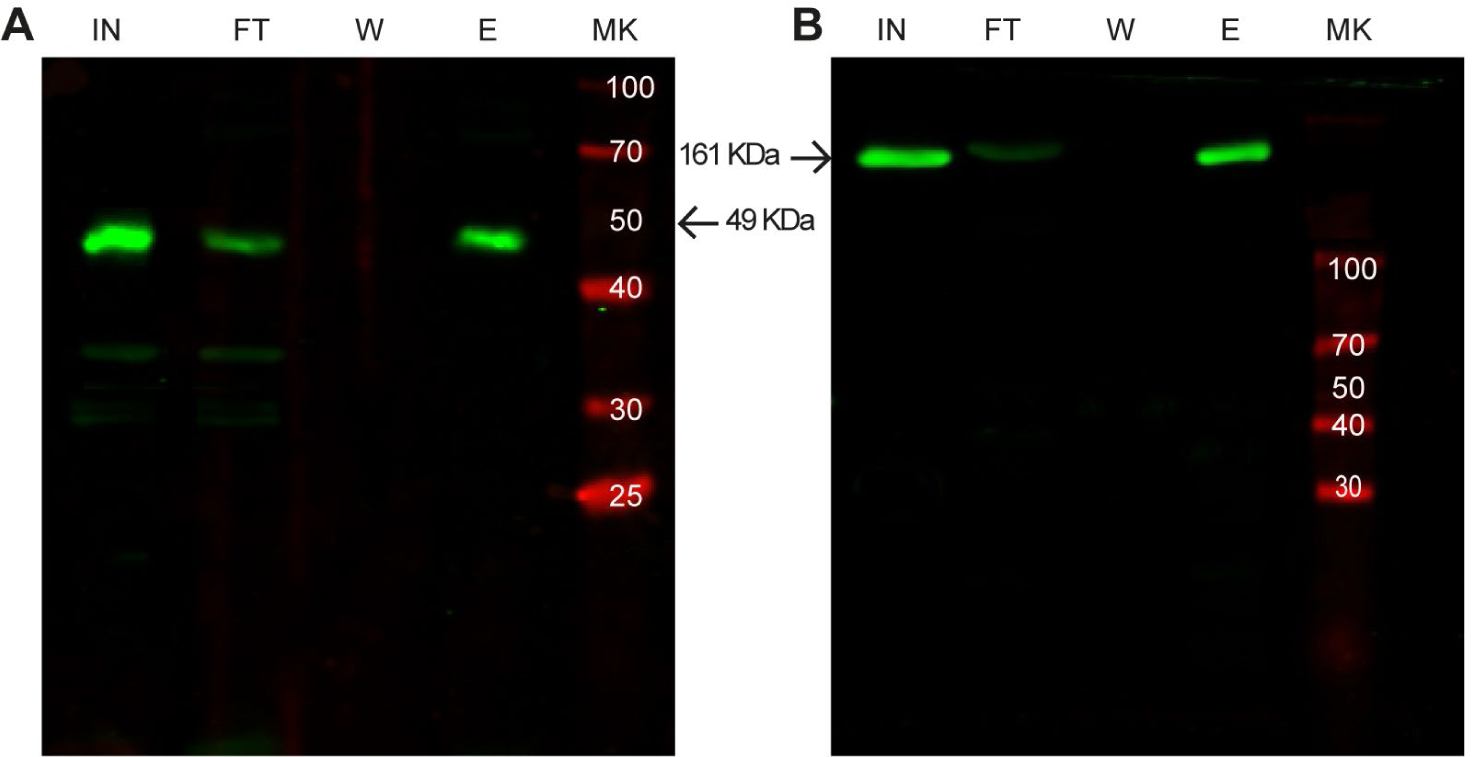
Western blot analysis of purification of SLAP-tagged proteins using different expression systems. (A) Purification of SLAP_ASE_ (human rhinovirus 3c fused to SLAP_TAG_) protease expressed in *Pichia pastoris*. (B) Purification of SARS CoV2 Spike-SLAP_TAG_ protein expressed in HEK293 cells. MK = protein marker (kDa); IN = input; FT = flowthrough; W = wash; E = elution.

### BM is a reusable chromatography matrix with long-term stability

As described in the Methods section a batch of BM was produced, and several aliquots were frozen at −20°C to study time stability. As shown in Fig. 7A, at different times, a few aliquots were unfrozen and tested for protein purification of the reporter protein (H_6_)-GFP-SLAP_TAG_ being the BM stable for more than a year that was tested (14 months). In addition, BM reusability capacity was evaluated determining that the matrix can be reused five times with no modification of the protein purification yield (Fig. 7B).

**Figure 7:**
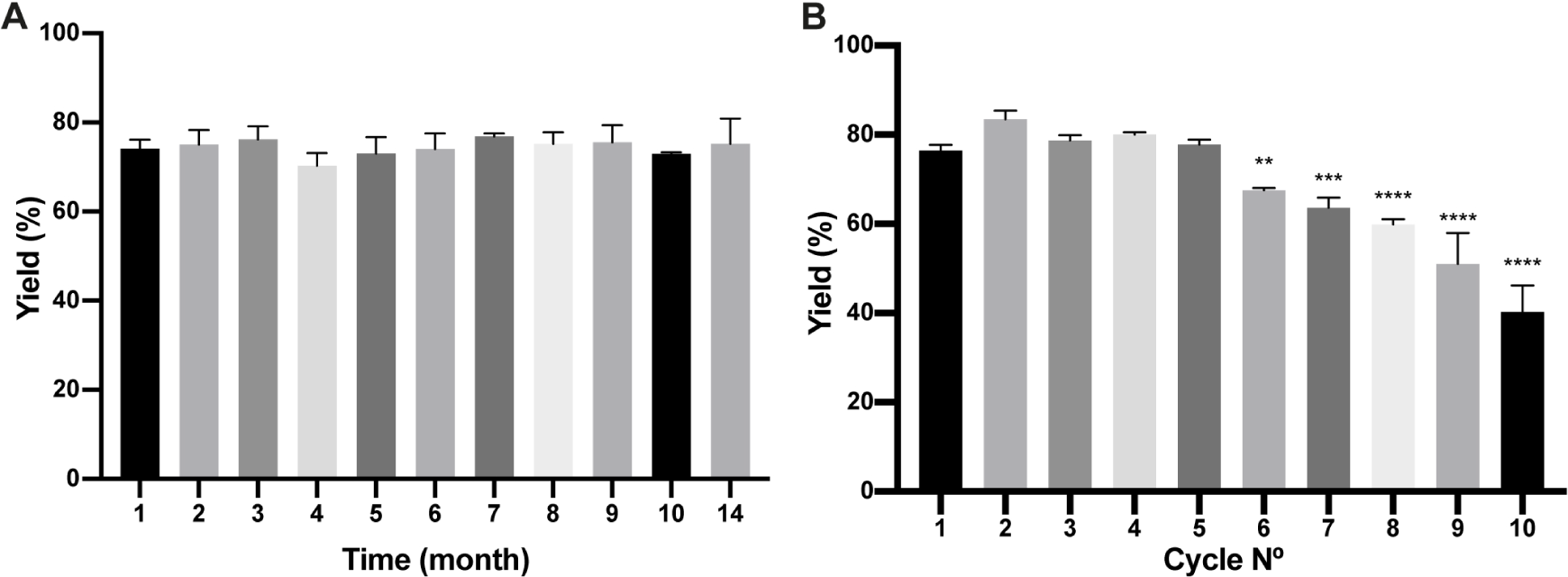
**Analysis of Bio-Matrix stability in time and reuse.** (A) The figure compares (H_6_)-GFP-SLAP_TAG_ purification yield of SLAP_TAG_-based affinity chromatography at different times when it is conserved at −20°C. Figure compares (H_6_)-GFP-SLAP_TAG_ purification yield of SLAP_TAG_-based affinity chromatography for different cycles of reuse. The asterisk denotes a significant difference using the ANOVA method, Bonferroni test (** p = 0,0059; *** p = 0,0002; **** p < 0.0001)

### Use of the Human Rhinovirus 3C protease fused to the SLAP_TAG_ (SLAP_ASE_) to release the SLAP-tagged proteins from BM

As shown in Fig. 4 and 5 the gene sequence of the viral protease HRV-3C was fused to the SLAP_TAG_ (named SLAP_ASE_), recombinantly expressed, and purified (Fig. 8B and 8D, lane SLAPase) to evaluate its activity. A reporter protein for SLAP_ASE_ activity was generated (GFP-_LEVLFQGP_-SLAP_TAG_) (Table 1) and a protocol for tag-removal by SLAPase was set as described in the Methods section. As shown in Fig. 8A the GFP-_LEVLFQGP_-SLAP_TAG_ was expressed (Fig 8B and Fig. 8C, lane Input or IN) and mixed with the BM for 5 minutes allowing the binding process. After binding, a purified SLAP_ASE_ was added to the mix and incubated for 60 minutes releasing a tag-less GFP (Fig. 8B and Fig 8C, Fig 8E, lanes flow-through or FT) by proteolysis. Cut SLAP_TAG_ is not observed in the flow-through indicating that was retained by the BM along with the SLAPase. To confirm these steps, post-proteolysis BM was incubated with LiCl solution to recover any residual bound SLAP-tagged protein. As shown in Fig. 8B and 8D (lane Elution or E), the LiCl solution released the SLAPase and the cut SLAP_TAG_.

**Figure 8:**
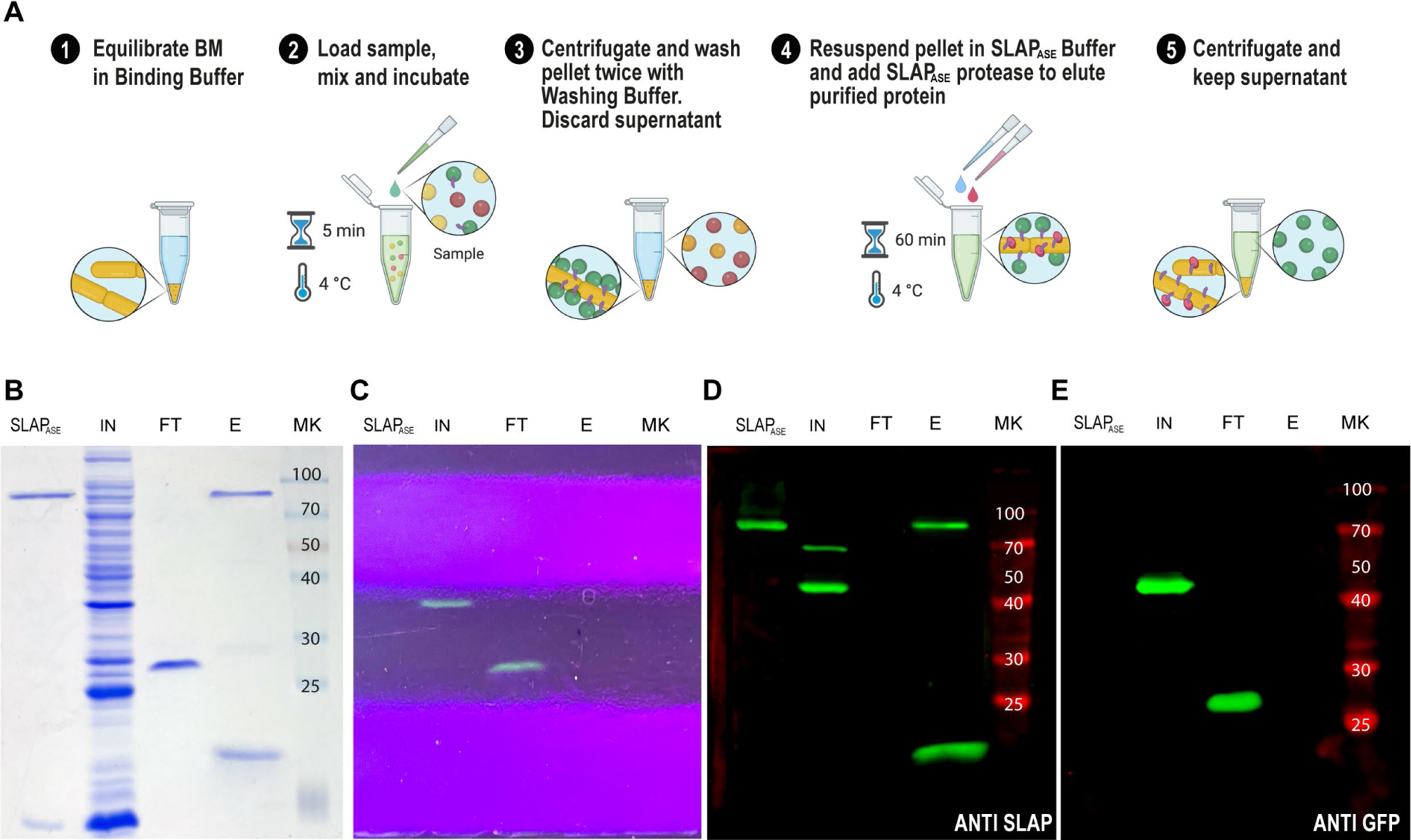
**SLAP_TAG_ removal.** (A) Graphical description of the protocol to remove the SLAP_TAG_ using SLAP_TAG_-based affinity chromatography. SDS-PAGE of purification fractions of SLAP_TAG_ removal protocol stained with Coomassie blue solution (B) or under UV light (C). Western Blot of purification fractions of SLAP_TAG_ removal protocol revealed with anti-SLAPTAG (D) or anti-GFP (E) antibodies. SLAP_ASE_ = purified protease; IN = input; FT = flowthrough; E = elution; MK = protein marker (KDa)

**TABLE 1:** Bacterial strains and plasmids used in this study

| Strain |  |  |
| --- | --- | --- |
| *E. coli* DH5α | F-φ80*lacZ*ΔM15 Δ(*lacZYA-argF*) U169 *recA1 endA1 hsdR17*(rK-, mK+) *phoA supE44λ- thi-1 gyrA96 relA1* | Invitrogen |
| *E. coli* BL21 Codon plus | [*ompT hsdS*(rB- mB-) *dcm* + Tcr *gal* λ (DE3) *endA* Hte Cmr] | Stratagen |
| *B. subtilis* | Wild type strain var. *natto* | ATCC 15245 |
| *Pichia pastoris* | *P. pastoris* strain GS115 | BioRad, USA |
| Plasmid |  |  |
| pET28-eGFP-SlpA | A vector containing the GFP gene from *Aequorea victoria* fused to SLAP<sub>TAG</sub> | 18 |
| pET22-eGFP-SlpA | A vector containing the GFP gene from *Aequorea victoria* fused to LEVLFQGP sequence and SLAP<sub>TAG</sub> | This study |
| pGEX-SlpA | A vector containing GST form *Shistosoma japonicum* fused to SLAP<sub>TAG</sub> | 12 |
| pLC3-EITH7-SlpA | A vector containing EspA-Intimin-Tir fused genes from STEC *E. coli* fused to SLAP<sub>TAG</sub> | 12 |
| pLC3-Omp19-SlpA | A vector containing the Omp19 gene from *Brucella abortus* fused to SLAP<sub>TAG</sub> | 12 |
| pLC3-FliC-SlpA | A vector containing the FliC gene from *Salmonella enterica* fused to SLAP<sub>TAG</sub> | 12 |
| pLC3-Gal8-SlpA | A vector containing the Gal8 gene from *Mus musculus* fused to SLAP<sub>TAG</sub> | 12 |
| pET28-Lys-SlpA | A vector containing the Lysozyme gene from *Gallus* fused to SLAP<sub>TAG</sub> | This study |
| pMAL-c5X-HRV3c-SlpA | A vector containing human rhinovirus (HRV) type 14 3C protease gene fused to SLAP<sub>TAG</sub> (SLAPase) | This study |
| pGEX-2T-EspA-SlpA | A vector containing the EspA gene from STEC *E. coli* fused to SLAP<sub>TAG</sub> | This study |
| pPICZalphaB-HRV3c-SlpA | A vector containing human rhinovirus (HRV) type 14 3C protease gene fused to SLAP<sub>TAG</sub> (SLAPase) | This study |
| Addgene #154754 | A vector containing the sequence of SARS CoV2 Spike ectodomain Hexa-pro | 29 |
| pSpike-SLAP<sub>TAG</sub> | Plasmid Addgene #154754 fused with the SLAP<sub>TAG</sub> | This study |
| pLysS | Vector for expression of T7 lysozyme. | Novagene |

### Adaptation of SAC to magnetic affinity chromatography

Since we confirmed the efficiency and universality of SAC for the purification of recombinant proteins, we explored a magnetic affinity chromatography alternative for SAC (Fig. 9A). As shown in Fig. 9B, 9C, and Supplementary Video-1 the (H_6_)-GFP-SLAP_TAG_ was purified in a few and easy steps. As shown in the SDS-PAGE (Fig. 9B) the adapted SAC protocol for magnetic chromatography (BM_mag_) was able to purify the reporter protein (H_6_)-GFP-SLAP_TAG_ very efficiently (Fig. 9B, lane Elution or E). Interestingly, this protocol can be adapted to commercial devices that use magnetic force for protein or DNA purification (like King Fisher Flex from ThermoFisher). In Fig. 9C the entire process of purification was monitored by direct observation under UV light (Supp. video 1). Noteworthy, the binding of the BM_mag_ to GFP quenched the fluorescence of this protein, an effect that was described for transition metal binding to GFP or binding of iron cations to fluorescent proteins (Fig. 9C, tube IN+BM_mag_) ^19, 20^.

**Figure 9:**
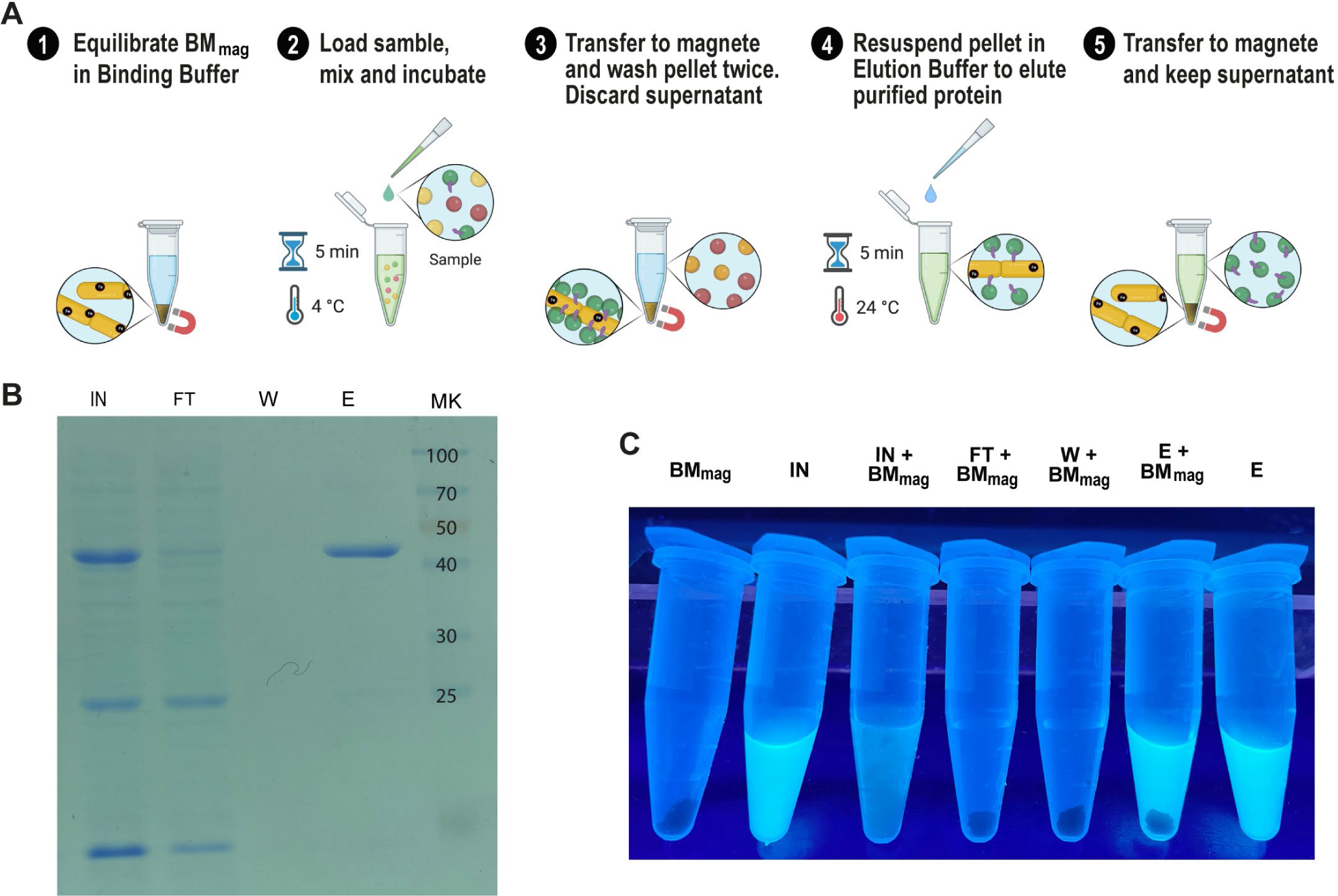
**GFP-SLAP_TAG_ purification using Magnetic Bio-Matrix.** (A) Graphical description of the purification protocol for magnetic Bio-Matrix. (B) Photo of 1,5 tubes ml containing fractions of GFP-SLAP_TAG_ magnetic purification under UV light. (BM_mag_ = magnetic Bio-Matrix; IN = input; FT = flowthrough; W = wash; E = elution. (C) Tubes containing magnetic purification fractions are seen under UV light. GFP input fluorescence is recovered in the eluate.

## Discussion

The *B. subtilis-derived* matrix, named here as Bio-Matrix (BM), showed a high performance in its binding and elution capacities, combined with good purification parameters for our reporter protein (H_6_)-GFP-SLAP_TAG_. Results presented here show that the SAC protocol can be potentially adapted as a universal tool for recombinant protein purification. Thus, proteins from different origins, with different molecular weights, or produced by different recombinant expression systems (bacteria, yeast, or cells) can be efficiently purified by SAC. In addition to its universal application, we demonstrate here that SAC was able to achieve a protein yield similar to those obtained by commercial affinity chromatography systems such as the Immobilized Metal Affinity Chromatography or IMAC. As reported, in protein production bioprocess, chromatography is the most expensive step^21^. Of interest, the SAC protocol only uses simple and low-cost reagents, consequently presenting an economic advantage in protein purification over current commercial systems. In addition, since the Biomatrix is “grown” instead of being chemically synthesized, the production of larger quantities of chromatography matrix can be achieved easily by simply scaling up the volume of the bioreactor. In addition, small research labs can easily produce their version of the in-house of BM by the protocol provided here.

One critical step in the generation of the affinity matrix is the immobilization of the affinity ligand to the chromatography matrix. Initially, at the beginning of the affinity chromatography development, the immobilization of affinity ligands was performed by covalent modification using diazo coupling ^11^. This procedure allows to immobilize different haptens or certain proteins to isolate antibodies. A second breakthrough in affinity chromatography was the development of the cyanogen bromide (CNBr) immobilization method which allows the easy cross-link of proteins or peptides to the activated agarose matrix^11^. These two major advances were combined in 1969 by Cuatreacasas et al where the term affinity chromatography was used for the first time ^22^.

In contrast, in Bio-Matrix affinity ligands (LTA and teichoic acid) are naturally integrated into the surface of the BM ^17^ and consequently, no chemical reactions to cross-link the affinity ligands to BM are required. Remarkably, no toxic chemicals or solvents are required in BM production.

From all the supports used as a matrix in affinity chromatography ^11^, the polysaccharide agarose is the most frequently used due to its low cost, large pores size, very low non-specific binding, and great stability over a broad pH range. However, agarose has limited mechanical stability at high operating pressures that limit the use of this matrix as support in high-performance liquid chromatography (HPLC) ^11^. Initially, other polysaccharides have also been adapted as supports in affinity chromatography like cellulose. Although cellulose displays a lower surface area and mechanical stability than agarose because of its low back pressure is a very useful tool used in preparative chromatography at high flow rates when used in membrane-based affinity separations^11^. In addition, carbohydrate-based supports combined with high-dense core material such as quartz, have been adapted in expanded-bed adsorption (EBA) chromatography^11^. Expanded-bed adsorption (EBA) chromatography is an excellent option for the capture of proteins directly from unclarified crude samples (such as a CHO cell supernatant)^23^. In EBA, the chromatography matrix (chromatographic bed) is initially expanded by an upward flow of the equilibration buffer^24^. A crude sample, that might have a mixture of soluble proteins, contaminants, entire cells, and cell debris along with the protein of interest (POI) is then passed upward through the expanded bed. In this process, POIs are captured on the adsorbent, while undesired particulates, aggregates, and contaminants pass through ^24^. Then, the replacement of binding buffer/washing buffer by an upward flow of elution buffer will result in POI desorption. Finally, when the flow is reversed, the adsorbent particle can be separated by sedimentation and desorbed POI can be recovered for further purification steps^24^.

EBA has been demonstrated to be a useful method, particularly for protein capture in a continuous protein purification process from unclarified feedstocks^25^ in the recovery of enzymes and therapeutic proteins from a variety of expression hosts^25^. Interestingly, the use of liquid magnetically stabilized fluidized beds (MSFBs) has been explored for EBA ^26^. Magnetically susceptible chromatography supports are forced to low back-mixing by applying a weak, external magnetic field that oriented the magnetic particles axially or transversely relative to the flow ^25^. This technique also gives great opportunities for process integration by achieving particulate removal and the capture of the product desired in a single operation^25 26^.

As shown here, SAC is an adsorption affinity chromatography, that was able to efficiently capture SLAP-tagged proteins directly from different feeds like bacterial lysates, *Pichia pastoris* supernatant, or HEK293 cells supernatant in a single step. In addition, a magnetically susceptible biomatrix was generated that still was able to capture SLAP-tagged proteins. Both results made SAC an interesting technique for EBA or EBA/MSFBs adaptation.

In the last decade, the introduction of single-use technologies has enlightened the potential for reduced regulatory and operational costs associated with chromatography. The researchers point to its potential for simpler operation, shorter processing times, and decreased buffer consumption, leading to better economics. ^27^. Also, the lack of need for cleaning over repeat-use cycles significantly reduces costs. SAC could be a single-use alternative chromatography for some industries, with the benefit of being more eco-friendly than those available on the market, as it is a biologically based and biodegradable matrix.

SAC proved to successfully adapt to protease tag removal. Although new technologies are being developed for tag removal (e.g., inteins), the enzymatic cleavage of the tag is still preferred as it is the most controlled process, with no premature cleaving and the best yields are obtained^28^. Moreover, new technologies might be compatible with SAC.

Therefore, we propose that SLAP_TAG_ affinity chromatography for protein purification in industries with permissive regulations. SAC can be adapted as an in-house protein purification system, available for any laboratory around the globe, to produce pure recombinant proteins for research, diagnosis, and the food industry. Although so far regulatory issues might preclude the use of the SAC for proteins used as biotherapeutics, new efforts have been performed to develop a new version of matrix chromatography suitable for more stringent industrial or clinical purposes.

## Methods

### Strains and plasmids

All the bacterial strains and plasmids used here were summarized in Table 1. Bacteria *Escherichia coli* and *Bacillus subtilis* strains were grown in Luria Bertani (LB) medium (Sigma, St. Louis, MO, United States) at 37°C and 180 rpm. Bacterial plasmid vectors were transformed into *E. coli* DH5α or *E. coli* BL21 (DE3) for protein expression. *Pichia pastoris* was grown in Yeast Extract–Peptone–Dextrose (YPD) medium at 28°C 180 rpm. HEK293F cells were maintained at 37°C in a 5% CO2 atmosphere in Dulbecco modified Eagle medium (DMEM) supplemented with 5% fetal bovine serum and streptomycin (50 μg/ml)-penicillin (50 U/ml). The SARS-CoV-2 Spike ectodomain Hexa-pro construction (Table 1) was a gift from Jason McLellan (Addgene # 154754; http://n2t.net/addgene:154754;RRID: addgene_154754)^29^.

### Cloning

Lysozyme gene was amplified from the pLysS plasmid (Millipore Sigma-Novagen) with the oligonucleotide primers Fw-lys-*Nde*I (CCCATATGGCTCGTGTACAGTTTAAACAACGTG) and Rv-lys-SalI (CGGTCGACTCCACGGTCAGAAGTGACCAGTTCG). The PCR product was digested with the restriction enzymes *Nde*I and *Sal*I and then cloned into the pLC3-EITH7-SlpA vector in the same restriction sites and the recombinant plasmid was transformed into *E. coli* DH5α. In addition, the lysozyme gene was subcloned into pET28-e GFP-SlpA at the *Nde*I and *Not*I restriction sites and transformed into *E. coli* BL21 by electroporation.

*EspA* gene was amplified with the following primers: *pRvEspAXbaI* (GCTCTAGATTTACCAAGGGATATTGCTG) *and pFwEITSalI* (ACGCGTCGACGATATGAATGAGGCATCTAAA). After digestion with *Xba*I and *Sal*I, the gene was cloned into *pGEX-SlpA* (Table 1), which was previously digested with the same enzymes. The plasmid was introduced by electroporation into *E. coli* BL21.

The *have-3c-slap tag* gene was synthesized and cloned in pMAL-c5X and pPICZalfa-B for protein expression in *E. coli* and *P. pastoris* respectively (Gene Universal, Delaware-USA).

For *P. pastoris* GS115 transformation, 5µg of pPICZalfa B – *hrv-3c* plasmid was linearized with *Sac*I restriction enzyme and transformed into cells through electroporation using 2 mm gap cuvettes (1500 V, 125 X, 50 lF). Transformed cells were selected by plating on a YPD medium containing Zeocin 100ug/ml for resistance selection. Isolated colonies were further grown in test tubes containing YPD broth and after 24 h, protein expression was induced by adding 1% (v/v) pure methanol every 24 h. Supernatants were sampled after 72 h of induction and the best producer clones were chosen for further experiments.

The pET28-eGFP-SLAP was digested with *BamH*I and *Xho*I restriction enzymes and the SLAP_TAG_-containing band was subcloned into SARS-CoV-2 S HexaPro plasmid (Addgene#154754) with the same restriction sites.

### Protein expression

*E. coli* BL21 were grown at 37°C 180 rpm in liquid LB supplemented with antibiotics until they reached OD_600_ = 0.6. Then, 0.1 mM IPTG was added, and bacteria were grown for another 20 hours at 18°C 180 rpm. Finally, bacteria were harvested and lyzed by sonication. The bacterial lysates were clarified by centrifugation.

*For Pichia, pastoris* protein expression cells were induced with methanol. Briefly, *P. pastoris* cultures were grown in 50 ml of YPD for 48 h. until the dextrose was consumed. Then, methanol pulses (200ul) were supplied every 24 h for 5 days. Finally, *P. pastoris* were harvested, and supernatants were collected.

HEK293 cells were transfected using polyethyleneimine (PEI) for protein expression. Briefly, thirty thousand cells per well were seeded in 24 well plates and incubated at 37 ℃, 5% CO_2_ for 24 h. For transfection, 4µl of PEI was diluted in 40µl of DMEM. 500 ng of DNA was added, and the transfection mix was incubated for 20 min at room temperature. Then, the transfection mix was transferred to the cells and incubated for 48 h. Finally, the supernatant was collected to check protein expression.

### 6xHis-Tag purification method

Purification of 6xHis-tagged proteins was carried out according to the manufacturer’s protocol. Briefly, benchtop columns were equilibrated with the binding buffer (200 mM NaCl, 50 mM Tris-HCl pH 7.5). Columns were loaded with the bacterial lysate. After being washed, columns were eluted in steps with 10, 50-, 100-, 300-, and 500-mM imidazole in the equilibration buffer. When compared with the SAC, the IMAC batch format was adopted. Thus, Ni-NTA superflow resin (Qiagen) was equilibrated with buffer (200 mM NaCl, 50 mM Tris-HCl pH 7.5) in 1,5 ml tubes. Cleared bacterial lysates were loaded and mixed in an orbital shaker for 1 hour at 4°C. Samples were centrifugated and, after being washed, the resin was eluted with 500 mM imidazole in an equilibration buffer.

### Bio-Matrix preparation

*B. subtilis* natto was grown in 200 mL of Luria-Bertani broth at 37°C 180 rpm for 48 hrs. Then, the culture was centrifugated and the bacteria were washed twice with PBS. The culture was resuspended in PBS with 2% glutaraldehyde and incubated overnight with soft agitation. Next, fixed bacteria were washed twice with PBS and stored in 20 % ethanol. 1mL of BM correspond to DO600=30 of B. subtilis.

BM was weighted in a drying scale (KERN MLS-D) and a calibration curve for dry weight vs optical density was performed.

### SLAP_TAG_ purification method

To purify the SLAP-tagged proteins, BM was equilibrated in binding buffer (200 mM NaCl, 50 mM Tris-HCl pH 7.5). Then, samples were incubated with the BM at different times and temperatures. Samples were centrifuged (8000 rpm) and the BM was washed 3 times with binding buffer. Finally, samples were incubated at different times and temperatures with the corresponding solution for elution. Proteins expressed in HEK293 and *P. pastoris* were purified from the supernatant with the final protocol described in this paper.

### Biomatrix time stability and reusability

To analyze stability in time, aliquots of BM were frozen at −20°C. At different times, samples were unfrozen and used for purifying GFP-SLAP following the protocol developed in this paper. To analyze reusage, an aliquot of BM was used for purifying GFP-SLAP following the protocol developed in this paper. After elution, the BM was washed with two volumes of elution Buffer (Bicarbonate Buffer pH 10, 200 mM NaCl) and then two volumes of binding buffer (200 mM NaCl, 50 mM Tris-HCl pH 7.5). The cleaning process was repeated each time after elution.

### Protein analysis

Protein samples were dissolved in a cracking buffer and incubated for 5 min at 100°C. Protein electrophoresis was performed at 120V on 12% SDS-PAGE gel. Gels were stained in a Coomassie-Blue solution (20% methanol, 10% acetic acid).

For Western Blot analysis, proteins were transferred to a nitrocellulose membrane for 55 min at 15V using a semi-dry electroblotting transfer unit (Bio-Rad, Hercules, CA, United States). Membranes were incubated for 1 hour with blocking buffer (1% dry skim milk, 0,1% Tween in PBS). Then, membranes were incubated for 1 hour with primary antibody diluted in blocking buffer (1/500). After washing with PBS-0,1% Tween, membranes were incubated for 1 hour with IRDye fluorophore-labeled secondary antibodies (LI-COR, Lincoln, NE, United States) diluted in a blocking buffer (1/20000). Finally, membranes were scanned using the Odyssey Imaging System (LI-COR).

### Fluorescence measurements

GFP fluorescence measurements were performed at 485/535nm excitation-emission wavelength respectively using FilterMax F5 Microplate Reader in Black 96 Well Plates (Thermo).

### Protein modeling

(H_6_)-GFP-SLAP_TAG_ structure was predicted by AlphaFold^13^. ColabFold web interface was employed using standard settings (five models and no templates).

### Adsorption isotherm

Adsorption isotherms for (H_6_)-GFP-SLAP_TAG_ on BM were performed using batch experiments. BM was equilibrated with binding buffer (50 mM Tris-HCl, 200 mM NaCl, and pH 7.6). Purified (H_6_)-GFP-SLAP_TAG_ protein at 3 mg/ml in binding buffer was used as a stock solution. Different concentrations of protein were incubated with 10 µl of BM. After reaching equilibrium, samples were centrifugated and GFP fluorescence from the supernatant was measured. Unbound protein in equilibrium with the BM was calculated using GFP fluorescence. Bound protein was estimated by the difference between input and unbound protein. The adsorption isotherm data were then fitted to the Langmuir isotherm equation to calculate the parameters Qmax and K_D_.

### Confocal Fluorescence Microscopy

Samples were incubated for 30 min in plates treated with poly-L-Lysine (Sigma, St. Louis, MO, United State). After treatment with PFA (4% in PBS), samples were washed twice with PBS. Finally, samples were observed with a confocal laser-scanning microscope Olympus FV1000 using a PlanApo N (60 × 1.42 NA) oil objective.

### Cleavage of SLAP_TAG_ with SLAP_ASE_ protease

Cleavage buffer recommended for commercial HRV3c protease was prepared: 50 mM Tris-HCl, pH 7.0, 150 mM NaCl, 1 mM EDTA, and 1 mM dithiothreitol. GFP-_LEVLFQGP_-SLAP_TAG_ protein was bound to the BM. After binding BM was washed with the same buffer thrice at 4°C. SLAP_ASE_ was added and incubated at 4°C. Percolate containing GFP protein without SLAP_TAG_ was recovered. As a control, the protease and SLAP_TAG_ were eluted with Bio-Matrix elution buffer after cleavage.

### Synthesis of iron nanoparticles

Iron nanoparticles were synthesized by reverse co-precipitation as described by Nadi et al ^30^. In summary, the precursor was prepared by dissolving 0.89 g of FeCl2.4H2O in 90 mL of water. The sample was incubated in stirring for 15 min and sonicated for 10 min to ensure complete dissolution of the salt. The solution was then poured over a solution of ammonium hydroxide diluted 1:2 with water and stirred for an hour. Finally, the sample was washed repeatedly with deionized water.

### Magnetic Bio-Matrix generation

Briefly, 5 ml of Bio-Matrix were resuspended in PBS and iron nanoparticles were added at a final concentration of 40 g/L. The sample was mixed with gentle stirring for 30 minutes. Then, it was washed repeatedly with PBS 1X and preserved in 20% ethanol.

### Antibody generation

Mice were intraperitoneally injected with 10µg of purified GST-SLAP_TAG_ or GFP in aluminum hydroxide (Imject Alum ThermoFisher). Two and four weeks later, booster doses of 5µg of protein were administrated.

### Statistical analysis

Statistical analyses were performed using GraphPad Prism 9 software. Statistical significance was analyzed by one-way ANOVA with Bonferroni.

## Acknowledgments

We would like to thank Dr. Juan E. Ugalde for his careful and critical reading of this manuscript.

This work was supported by grants from the Agencia Nacional de Promoción Científica y Técnológica, Buenos Aires, Argentina, PICT-2018-0778, PICT-2021-CAT-I-00049 and CONICET (PUE-0086). E.J.M., P.J.U., and C. M. D. are doctoral fellows from CONICET. MVP is a postdoctoral fellow from the Agencia Nacional de Promoción Científica y Tecnológica. M.S.R. and G.B. are members of the Research Career of CONICET.

## Availability of materials and data

All data generated or analyzed during this study are included in this published article and its supplementary information files.

---

**Table S1:** Purification table of GFP-SLAP_TAG_ using the Biomatrix

**Supplementary Video 1:** Magnetic affinity chromatography using BIOMATRIX

