## Supplementary material for "Adaptation of the binding domain of *Lactobacillus acidophilus* S-layer protein as a molecular tag for affinity chromatography development": Purification table (from another version of this preprint: version 1, 2022-12-25)

Table S1: Purification table of GFP-SLAP_TAG_ using the biomatrix

| Step | Vol (ml) | Total protein (mg) | Total activity (RFU*) | Specific Activity (RFU/mg) | Purification fold | Yield (%) | Efficiency |
| --- | --- | --- | --- | --- | --- | --- | --- |
| Crude extract | 10 | 591 | 3,75 x10^11^ | 6,34 x10^8^ | 1 | 100 | - |
| Biomatrix | 1 | 5,96 ± 0,30 | 2,96x10^11^ ± 1,57 x10^10^ | 4,96 x10^10^ ± 2,92 x10^9^ | 78,23 ± 4,61 | 78,90 ± 4,20 | 61,81 ± 6,33 |

***Activity RFU (relative fluorescence units)**
